# Learning-induced reorganization of neuronal subnetworks in the primary sensory cortex

**DOI:** 10.1101/2023.02.21.529414

**Authors:** Yexin Yang, Hao Shen, Sung Eun Kwon

## Abstract

Perceptual learning alters the representation of sensory input in primary sensory cortex. Alterations in neuronal tuning, correlation structure and population activity across many subcortical and cortical areas have been observed in previous studies. However, relationships between these different neural correlates - and to what extent they are relevant to specific perceptual tasks - are still unclear. In this study, we recorded activity of the layer 2/3 neuronal populations in the whisker primary somatosensory cortex (wS1) using in vivo two-photon calcium imaging as mice were trained to perform a self-initiated, whisker vibration frequency discrimination task. Individual wS1 neurons displayed learning-induced broadening of frequency sensitivity within task-related categories only during task performance, reflecting both learning-and context-dependent enhancement of category selectivity. Learning increased both signal and noise correlations within pairs of neurons that prefer the same stimulus category (‘within-pool’), whereas learning decreased neuronal correlations between neuron pairs that prefer different categories (‘across-pool’). Increased noise correlations in trained animals resulted in less accurate decoding of stimulus categories from population activity but did not affect decoding of the animal’s decision to respond to stimuli. Importantly, within-pool noise correlations were elevated on trials in which animals generated the learned behavioral response. We demonstrate that learning drives formation of task-relevant ‘like-to-like’ layer 2/3 subnetworks in the primary sensory cortex that may facilitate execution of learned behavioral responses.

**Significance Statement:** We found that cortical plasticity during perceptual learning alters both neuronal tuning and the structure of pairwise correlations such that they become increasingly aligned to task-related categories, indicating the formation of ‘like-to-like’ subnetworks in layer 2/3 of sensory cortex. Category-specific increases in signal and noise correlations were induced by learning and only observed during active task performance, which points to top-down feedback as a driver of task-related subnetworks.

## Introduction

Neurons in primary sensory cortex encode sensory information that drives perception and behavior (Buetfering et al., 2022; Carrillo-Reid et al., 2019; Kwon et al., 2016; Marshel et al., 2019; Sachidhanandam et al., 2013; Yang et al., 2016). Representations of sensory input in primary sensory cortex remain plastic even in mature brains. Cortical plasticity is thought to underlie important cognitive processes like perceptual learning; training-induced perceptual improvements in discrimination or detection tasks that occur throughout life. The exact nature of plasticity processes and the impact of these changes on the neural representation of task-relevant sensory input remain debated. Prior studies in rodents report that perceptual learning induces alterations of neuronal tuning, correlation structure and/or population activity in early sensory cortical areas (Caras and Sanes, 2017; Corbo et al., 2022; Goltstein et al., 2021; Khan et al., 2018; Peron et al., 2015; Schumacher et al., 2022; Xin et al., 2019). Critically, however, relationships between these different neural correlates of learning - and to what extent they are relevant to specific perceptual tasks - are still unclear. To address these issues, we longitudinally characterized neuronal tuning, correlation structure and population coding in sensory cortex of animals learning a tactile frequency discrimination task. We quantified changes in these measures either before and after learning, or across repeated exposure to sensory stimuli, to extract learning-induced modifications in the cortical circuit.

We recorded activity of the layer 2/3 neuronal populations in the whisker primary somatosensory cortex (wS1) using *in vivo* two-photon calcium imaging as mice were trained to perform a self-initiated, whisker vibration frequency discrimination task. Individual wS1 neurons displayed learning-induced broadening of frequency sensitivity within stimulus categories related to the task (i.e., Go vs. NoGo), reflecting an enhanced category selectivity after learning. To test if such changes lead to network-level reorganization, we compared pairwise neuronal correlation matrices before and after training. We found that learning increased both signal and noise correlations within pairs of neurons that prefer the same stimulus category (‘within-pool’), whereas learning decreased neuronal correlations between neuron pairs that prefer different categories (‘across-pool’). Learning specifically induced alterations of neuronal correlations as passive exposure to the repeated whisker stimulation had little effect. Importantly, within-pool noise correlations were significantly elevated on trials in which animals generated the correct learned behavioral response. Increased noise correlations in trained animals resulted in less accurate decoding of stimulus category from population activity but did not alter decoding of the animal’s decision to respond to stimuli. Taken together, our results show that learning drives formation of task-relevant ‘like-to-like’ neuronal subnetworks in layer 2/3 (L2/3) of the primary sensory cortex. These subnetworks appear to facilitate execution of learned behavioral responses, despite their detrimental effect on encoding of sensory information.

## Materials and Methods

### EXPERIMENATL DESIGN

#### Mice

All procedures were in accordance with protocols approved by the University of Michigan Animal Care and Use Committee. We report calcium imaging experiments from 10 C57BL/J6 (Jackson Labs) mice from a C57BL/J6 background, with ages greater than 5 weeks (*SI Appendix*). Both sexes were used. Mice were housed in a vivarium with a reverse light-dark cycle (12 h each phase). Experiments occurred during the dark phase. After recovery from surgery (see below), mice were singly housed and water-restricted by giving them 1 mL per day. Mouse weight did not go below 70% of the starting weight.

#### Surgery and Virus Injection

Mice were anesthetized with 1% isoflurane throughout surgery and kept on a thermal blanket to maintain body temperature. The scalp and periosteum over the skull were carefully removed. A circular craniotomy was made on the left hemisphere (3.0 mm diameter) with the dura left intact. The center of the craniotomy was located over the wS1 barrel cortex (3.5 mm lateral and 1.3 mm caudal relative to Bregma). Injections were performed unilaterally using a beveled glass pipette (30-50 μm diameter) mounted on an oil-based hydraulic micromanipulator (Narishige). Adeno-associated virus for expressing GcaMP7f or 8f under the synapsin-1 promoter (AAV1-syn-jGCaMP7f-WPRE; Addgene 104488; AAV1-syn-jGCaMP7f-WPRE, Addgene 162376) was injected into the wS1 at depths of 250 μm and 120 μm below the dura and at a rate of 1 nL/sec (50 nL total per location). Injection was made at 3 different locations on the cortical surface around the coordinates given above. The craniotomy was covered with a glass window after the injection. The window was made by gluing two pieces of coverslip glass together. The smaller piece (3.0 mm diameter) was placed into the craniotomy and while the larger piece (4.0 mm diameter) was glued to the bone surrounding the craniotomy. Cyanoacrylate adhesive (KrazyGlue) and dental acrylic (Jet Repair Acrylic) were used to secure a titanium head post in place on the skull. Silicone elastomer (Kwik-Cast, WPI) was placed over the window for protection during the recovery period. The mouse was allowed to recover for at least 10 days before moving to water restriction. Imaging started 3-5 weeks after surgery.

#### Behavioral Task

Head-restrained mice were trained to perform a frequency discrimination task using a behavioral apparatus controlled by BPod (Sanworks). Mice were placed in an acrylic (4.5 cm inner diameter) tube. For 7–10 d before training, mice received 1 ml per day of water. Mice were weighed prior to and after training sessions to ensure the amount of water consumed. In the first 3 sessions (‘Habituation’), mice were allowed to freely lick at the water port positioned near its snout. Each time the tongue crossed the infrared beam to touch the water port, the mouse received a drop of water (~7 μL). In the next 2 sessions (‘Go only’), mice were conditioned to lick at the water port to a passive whisker deflection. Facial whiskers were threaded through a plastic mesh attached to a piezoelectric actuator (CTS). For training in ‘Go/NoGo’ sessions (2-4 days), the whiskers were deflected for 1 s with sinusoidal deflection (rostral to caudal) at 30 Hz (Go trials) or 5 Hz (NoGo trials). Go and NoGo trials each comprised 50% of all trials, therefore stimulus categories were equally represented in the data. 0.1-s auditory tone (8 kHz, ~70 dB SPL) was delivered starting 2 s before whisker stimulus onset, followed by a 0.5-s ‘No-lick’ window. If mice lick during this window, the trial was defined as ‘Premature Licking’ and aborted. Licks occurring during the first 1.2 sec after the onset of whisker deflection had no consequence. The ‘reward window’ was defined as 1.2 – 3.2 s after the onset of whisker deflection. The Go trials resulted in a ‘hit’ when the mouse licked the water port within the reward window and received a drop of water. A ‘miss’ occurred if mice did not lick within the reward window, and no reward or punishment was delivered. The NoGo trials resulted in a ‘false alarm’ if mice had licked within the reward window, and mice were punished by a 5.00 s time-out. Licking during time-out resulted in an additional time-out. A ‘correct rejection’ occurred if mice did not lick within the reward window on NoGo trials. In the next 2-4 sessions (‘Self-initiated Go/NoGo’), mice were trained to initiate each Go or NoGo trial by turning a wheel (Lego) attached to a rotary encoder more than 65 degree in either direction. The task itself was otherwise identical to the previous stage. At the last phase (‘discrimination task’), the whiskers were stimulated for 1 s at a varied frequency. In ‘High-Go’ paradigm, whiskers were deflected for 1 s at a frequency randomly selected from 20, 25, 30, 40, 50 Hz on ‘Go’ trials and 2, 5, 10, 15 Hz on ‘NoGo’ trials. In ‘Low-Go’ version of the task, the contingency was reversed. During all sessions, ambient white noise (cut off at 40 kHz, ~60 dB SPL) was played through a separate speaker to mask any other potential auditory cues associated with movement of the piezoelectric actuator. No more than 3 trials of the same type occurred in a row. The fraction of correct trials (Fraction Correct) was defined as the number of hits plus correct rejections divided by the total number of Go and NoGo trials. The hit rate was defined as the number of hits divided by the total number of Go trials. The false alarm rate was defined as the number of false alarms divided by the total number of NoGo trials. Mice were considered trained once the performance was > 70 % correct. d-prime was calculated as z(hit rate) - z(false alarm rate). Mice reached this performance criteria after 10−12 daily training sessions.

#### wS1 lesion and silencing experiments

After confirming that mice reached >70% performance in the frequency discrimination task, we inactivated wS1 by either aspirating the cortical tissue with gentle vacuum suction or stereotaxically injecting GABAa receptor agonist muscimol in the wS1 (3.5 mm lateral and 1.3 mm caudal relative to Bregma) under anesthesia with 1% isoflurane. For silencing with muscimol (5 mg / ml), injections occurred at three different depths (200, 400, 600 μm; 50 nl each) from the pial surface. The craniotomy was sealed with silicone sealant (KwikCast) following injection, and covered with a thin layer of dental cement. Behavioral performance in the frequency discrimination task was monitored after recovery from anesthesia (1 h). In the following day, we tested the effect of saline injected into wS1 on behavior using the same method as outlined above.

#### Two-Photon Calcium Imaging of Layer 2/3 Somata

Images were acquired on a Scientifica two-photon microscope (Hyperscope) equipped with an 8 kHz resonant scanning module, 2 GaAsP photomultiplier tube modules, and a 16× 0.8 NA microscope objective (Nikon). GCaMP was excited at 960 nm (40-60 mW at specimen) with an InSight X3 tunable ultrafast Ti:Sapphire laser (Spectra-Physics, Santa Clara, CA, USA). Imaging fields were restricted to areas where GCaMP expression overlapped with the center of the cranial window (3.5 mm lateral and 1.3 mm caudal to Bregma). The beam was focused to 150 – 250 μm from the cortical surface. The field of view ranged from 458 μm x 344 μm to 275 μm x 207 μm. Images were acquired with a resolution of 512 x 512 pixels at 30 Hz using ScanImage. A movie for a single trial consisted of 200 frames.

### STATISTICAL ANALYSIS

#### Image Analysis

After correcting for motion, regions of interest (ROIs) were selected and then manually curated to remove ROIs that were not neurons. The neuropil fluorescence time series was multiplied with a correction factor of 0.7 and then subtracted from the raw fluorescence time series to obtain the corrected fluorescence time series: Fcorrected(t) = Fraw - Fneuropil * 0.7. ΔF/Fo was calculated as (F-Fo)/Fo, where Fo represents the baseline fluorescence calculated by determining the average fluorescence (F) during 500-167 ms time window preceding whisker stimulus onset. Evoked ΔF/Fo responses were calculated as the average ΔF/Fo over 21 frames following the onset of whisker stimulus delivery minus the average ΔF/Fo over the 5-15th frames preceding the onset of whisker stimulus delivery. To assign each neuron as ‘responsive’ or ‘non-responsive’, the averaged standard deviation of baseline ΔF/Fo for each neuron was calculated. For each neuron each trial, if there are at least 3 frames during the 21 frames after onset of stimulus have ΔF/Fo three standard deviations greater than Fo, the neuron was considered responsive at that trial. Neurons that were responsive in more than 8% of the trials during the session were considered responsive neurons.

#### Single Neuron ROC Analysis

The receiver operating characteristic (ROC) analysis was used to calculate the ‘stimulus probability’. A decision variable (DV) was assigned for each trial based on the response of the particular neuron. DV was the evoked ΔF/Fo as defined above. Trials were grouped by the stimulus category (High vs. Low) and for each behavioral state for stimulus probability and an ROC curve was obtained by systematically varying the criterion value across the full range of DV using the MATLAB ‘perfcurve’ function. The area under the ROC curve (AUC) represents performance for an ideal observer in categorizing trials based on the DV. Stimulus probability was the AUC for discriminating the stimulus condition.

#### Noise and Signal Correlation Analyses

We calculated across-neuron pairwise noise correlations between neuron pairs recorded at the same time in a single session, across trials sharing the same stimulus frequency (Kwon et al., 2018). Evoked ΔF/Fo responses for each neuron were z-scored within each stimulus frequency. Noise correlation was then calculated as the Pearson correlation coefficient of the z-scored responses of two neurons across trials. We calculated pairwise signal correlations as the similarity of frequency tuning between neuron pairs recorded at the same time in a single session. Average evoked ΔF/Fo responses were calculated for each stimulus frequency and z-scored across all stimulus frequencies for each neuron. The stimulus frequency associated with maximal z-score was defined as the preferred frequency for each neuron. Signal correlation was calculated as the Pearson correlation coefficient of the z-scored responses of two neurons across stimulus frequencies. To cancel out the effect of noise correlation without impacting the tuning property of individual neuron, trial labels were shuffled 100 times within each stimulating frequency for each neuron.

#### Population Decoding

The Support Vector Machine (SVM, Python sklearn package) with linear kernel was used to decode the stimulus categories or behavioral choices from the vector of z-scored evoked ΔF/Fo responses. For each session, the SVM classifier was trained using labels for stimulus (1 for ‘Go’, 0 for “Nogo’) or behavioral choices (1 for ‘Licking’, 0 for ‘Nolick’) and tested with a nested 5-fold cross-validation scheme. 80% of the trials were randomly selected for training and the remaining 20% of the trials were used for testing. Classifier performance was evaluated as label-prediction accuracy. The final prediction accuracy was generated from 100 times of repeating training and testing classifier. All classifiers were regularized using L2-regularization. The regularization parameter was optimized through a pre-training session where the best penalty parameter that yields the highest accuracy was chosen. According to design of the behavioral paradigm, each session had well-balanced number of trials of two stimulus categories but may have unbalanced choices. To avoid the effect of unbalanced numbers of trials, subsets of ‘Lick’ trials or ‘No-lick’ trials were randomly subsampled to match the number of trials across different trial types.

#### Image Analysis

Image stacks were processed using Suite2P pipeline (Stringer and Pachitariu, 2019). Procedures for ROC analysis, neuronal correlation calculations and population decoding analysis are described in *SI Appendix, Extended Methods*.

#### Statistics

Error bars indicate mean ± SEM unless mentioned otherwise. All statistical tests are two-tailed.

## Results

### Mice learn to discriminate whisker vibration frequencies using wS1

We trained head-fixed mice to perform a vibration frequency discrimination task in which they report, by licking or withholding licking, whether whiskers received a high- or low-frequency sinusoidal deflection (**Figure 1A**). Mice initiated each trial by manually turning a wheel, which was followed by a brief auditory tone (8 kHz, 0.1 s, ~70 dB SPL) indicating the beginning of a trial. Facial whiskers were threaded into a mesh attached to a piezoelectric bender and deflected for 1 s at a frequency randomly selected from 20, 25, 30, 40, 50 Hz on ‘Go’ trials (50% of all trials). On ‘NoGo’ trials (50% of all trials), whiskers were deflected at a frequency selected from 2, 5, 10, 15 Hz. A response window where Go cued licking was rewarded began after a 1.2-s delay following the onset of whisker deflection. Licking during the delay period had no behavioral consequences. Licking to the NoGo cue during the response window resulted in a brief (3 s) time-out. Trial outcomes comprised a mixture of successful responses (“Hits”) and failed responses (“Misses”) following Go stimulus delivery, as well as correct omission of licking (“Correct Rejection”) and incorrect (“False Alarms”) licking responses in the presence of the NoGo stimulus. Premature licking during the ‘No-lick window’, which lasted for 0.5 s after the trial initiation, led to a trial abortion.

**Figure 1.**
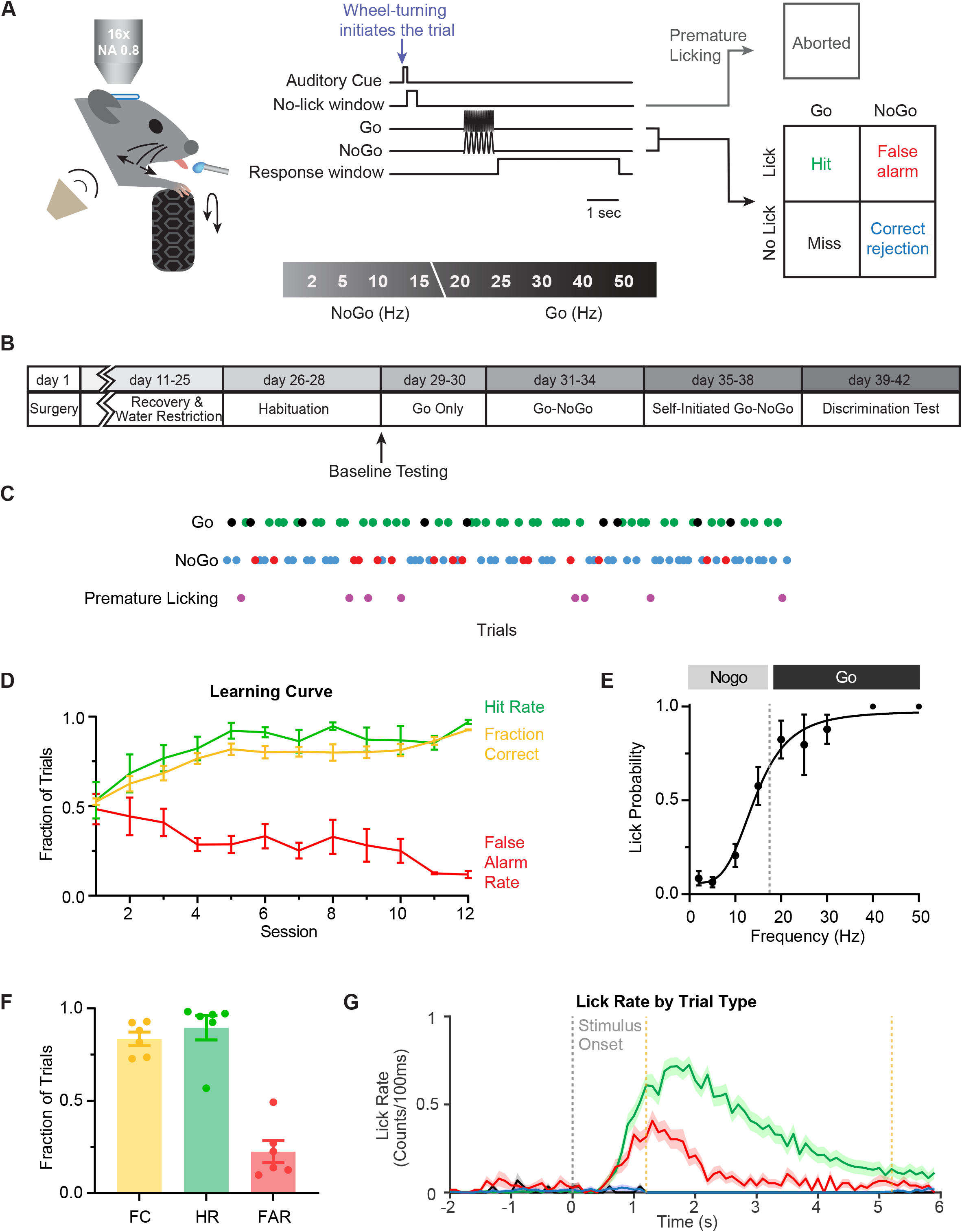
Whisker vibration frequency discrimination task in head-fixed mice. (A) Schematic showing behavior set-up and task structure. Water-restricted mice were trained to initiate each trial and lick the water port if the frequency of whisker vibration by a piezoelectric stimulator is ‘high’ (20, 25, 30, 40 or 50 Hz; Go cue) and withhold licking if it is ‘low’ (2, 5, 10 or 15 Hz; NoGo cue). Go and NoGo trials each comprised 50% of all trials. A 0.1-s auditory tone (8 kHz, ~70 dB SPL) was delivered starting 2 s before whisker stimulus onset, followed by a 0.5-s ‘No-lick’ window. If mice licked during this window, the trial was defined as ‘Premature Licking’ and aborted. Licks occurring during the first 1.2 sec after whisker stimulus onset had no consequence. The ‘reward window’ was defined as 1.2 – 5.2 s after the whisker stimulus onset. (B) Experimental timeline. (C) Representative session showing trials with hit (green), miss (black), false alarm (red), correct rejection (blue) and premature licking (purple). (D) Behavioral performance averaged across late lickers trained under ‘High-Go’ paradigm as a function of training sessions (n = 6 mice). (E) Average psychometric function of all animals trained under ‘High-Go’ paradigm (n = 6 mice). The dashed black line marks the experimentally defined Go/NoGo category boundary. (F) Average fraction of correct trials (FC), hit rate (HR) and false alarm rate (FAR) of expert mice (n = 6 mice). FC: 0.836 ± 0.037; HR: 0.896 ± 0.066; FAR: 0.225 ± 0.060. (G) Distribution of licks in 100 ms bins within trials with hit (green), miss (black), false alarm (red), correct rejection (blue). The dashed gray line indicates the onset of whisker deflection. The dashed yellow lines mark the onset and offset of the reward window.

Mice were trained in multiple phases (**Figure 1B**). After habituation to head-fixation, they performed the ‘Go Only’ task which included trials with 30 Hz whisker vibration only. Licking during the response window was rewarded. During the ‘Go / NoGo’ phase, mice were trained to discriminate between 30 Hz (‘Go’) and 5 Hz (‘NoGo’) stimuli. When the fraction of correct trials reached 80%, the task was switched to ‘Self-initiated Go / NoGo’ phase where they learned to initiate each trial by manually turning a wheel. Finally, in the ‘Discrimination Test’ phase, the full range of vibration frequencies were introduced. Mice typically completed 120-150 trials per session (**Figure 1C**). A separate group of mice was subjected to a ‘Passive Exposure’ condition. These mice were water-restricted and went through the same number of sessions as the experimental group except that licking had no behavioral consequences under this condition. They were allowed to drink water freely from the lick-port after each training session ended.

The fraction of correct trials increased throughout training, mainly due to a gradual decrease in the fraction of incorrect responses on NoGo trials (False Alarm Rate) (**Figure 1D**). It typically took 10-12 daily training sessions for mice to become an ‘expert’ at the discrimination task, defined as the fraction of correct trials greater than 0.7. The performance curve of expert mice shows a sigmoid relationship between lick probability and stimulus frequency, indicating categorical decision making (**Figure 1E**). Expert mice licked preferentially in response to Go cues (fraction correct: 0.836 ± 0.037; hit rate: 0.896 ± 0.066; false alarm rate: 0.225 ± 0.060; n = 6 mice) (**Figure 1F**). Lack of response is somewhat difficult to interpret in a typical Go / NoGo task. The self-paced nature of this task resolves this issue to some extent; not licking reflects a correct behavioral response (on NoGo trials) or a perceptual error (on Go trials) rather than inattention or lack of motivation. After training, number of licks on Go trials did not peak immediately, but with a delay of ~1.5 s after the stimulus onset, probably due to the 1.2 s delay before the beginning of response window. Mice consistently withheld licking during the first 0.70 s of the 1.2 s delay period (**Figure 1G**).

Before characterizing learning-induced modifications in wS1, we tested if wS1 contributes to the tactile frequency discrimination. We inactivated wS1 by aspirating the cortical tissue (3 mice) or locally infusing the GABA_A_ agonist muscimol (1 mouse) (**Figure 2A**). After tissue aspiration or muscimol infusion, the slope of the performance curve decreased in all mice, compared with pre-lesion performance (**Figures 2B and 2C**). Performance curves of individual mice reflected the same trend (**Figures 2D**). This result is consistent with previous studies showing that acute inactivation of the wS1 impairs the task performance (Hong et al., 2018; Miyashita and Feldman, 2013). Licking on NoGo cues, including in response to 5 Hz stimuli, increased with wS1 inactivation but returned to baseline level with additional training, consistent with previous studies (**Figure 2E**) (Hong et al., 2018; Hutson and Masterton, 1986). These results demonstrate that wS1 is critical for efficient tactile frequency discrimination.

**Figure 2.**
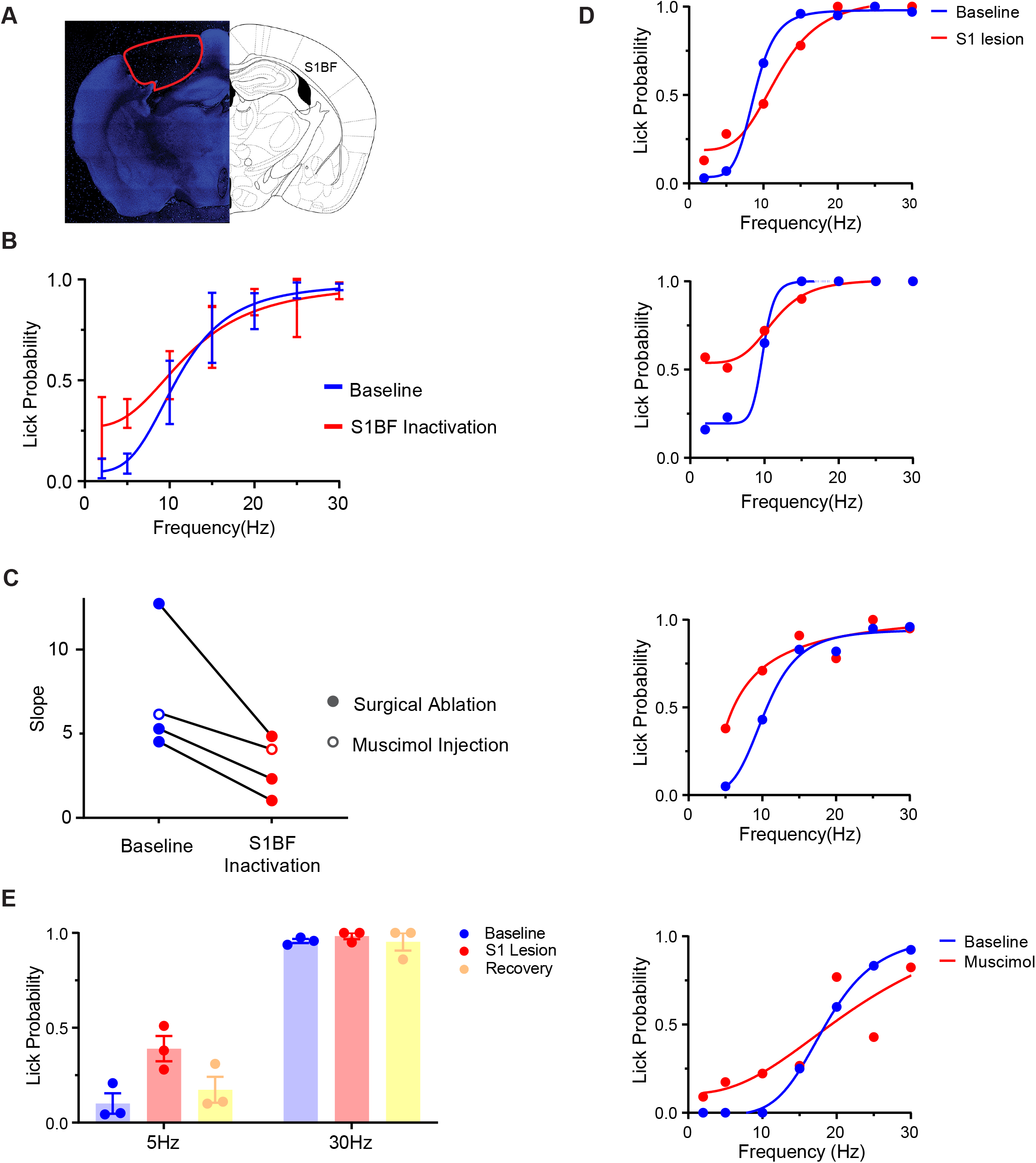
Efficient tactile frequency discrimination requires intact wS1. (A) wS1 containing barrel field (BF) was carefully removed in animals trained to perform the frequency discrimination task. Red line outlines the cortical tissue that was removed. (B) Psychometric curves that depict the probability with which the stimulus was perceived ‘high’ in relation to stimulus frequency, before versus after the wS1 inactivation. (C) The slope of psychometric function was reduced in all tested animals (n = 4 mice). (D1-3) Performance curves of well-trained individual mice at baseline (blue) and 1 day post wS1 removal (red). (D4) Performance curve of a well-trained mouse at baseline (blue) and on the day of wS1 muscimol injection (red). (E) Average lick probability on 5 Hz and 30 Hz trials at baseline (blue), 1 day (red) and 4 days (yellow) post wS1 removal (n = 3 mice).

### Learning enhances category representation among L2/3 neurons of wS1

To determine the neural correlates of learning, we monitored stimulus-evoked responses of L2/3 neurons using *in vivo* two-photon imaging of the genetically encoded calcium indicator GCaMP7f or 8f (**Figure 3A**) (Dana et al., 2019). We compared the frequency selectivity of individual neurons in response to whisker vibration at nine different frequencies randomly delivered during task performance before and after mice learned the task. Our analysis was based on comparisons of the same imaging field, and the same subpopulation of neurons, before versus after learning (**Figure 3B**), although responses from individual neurons were not compared across training sessions. To characterize frequency tuning of individual neurons, mean stimulus-evoked *ΔF/F_0_* responses were calculated using frames within 0.7 s (21 frames) after whisker stimulus onset, and they were z-scored across all frequencies. This time window was used in all subsequent analyses to avoid potential confounding effects related to licking (**Figure 1G**). The stimulus frequency that elicited maximal z-scored *ΔF/F_0_* in a given neuron was defined as its ‘preferred frequency’. We pooled neuronal tuning curves (i.e., z-scored *ΔF/F_0_* across nine different frequencies) across mice and sorted them by their preferred frequency. All tested vibration frequencies were represented by L2/3 neurons in wS1 of naive mice (**Figures 3C and 3D**). Importantly, training in the frequency discrimination task resulted in broadening of neuronal tuning curves within Go and NoGo stimulus categories (**Figure 3C**), whereas passive exposure had little effect on tuning curves (**Figure 3D**). We averaged the nine *ΔF/F_0_* z-scores (i.e. individual tuning curves) across subpopulations of neurons that share the same preferred frequency. Learning resulted in broadening of subpopulation tuning curves at most frequencies within each category (NoGo: 2, 5 and 10 Hz; Go: 30, 40 and 50 Hz), whereas they remained relatively unchanged at categorical boundary frequencies (15 Hz and 20 Hz) (**Figure 3E**). We also compared the frequency tuning curves before versus after passive exposure in a separate group of mice. Little change in tuning curves was observed under this passive exposure condition (**Figure 3F**). To gain insight into learning-induced changes in the category selectivity of individual neurons, we quantified the response selectivity for Go versus NoGo category using receiver operating characteristic (ROC) analysis. ‘Stimulus Probability’ captures how well an ideal observer could categorize a sensory stimulus (in our case, high vs. low-frequency vibration) based on the neural response in a trial-by-trial manner with 0.5 indicating lack of selectivity towards either category (Kwon et al., 2016). We used stimulus-evoked *△F/F_o_* as a decision variable for individual trials. Training in the discrimination task pushed the distribution of stimulus probability towards the two extremes (median [IQR] for pre-training: 0.495 [0.098]; post-training: 0.459 [0.331]), indicating an increase in the number of neurons representing either stimulus category (**Figure 4A**). Passive exposure to repeated whisker stimulation also slightly increased the width of stimulus probability distribution (**Figure 4B**), but the effect was subtle compared with active discrimination learning (median [IQR] for pre-training: 0.528 [0.081]; post-training: 0.508 [0.133]). Therefore, tactile discrimination learning enhances encoding of stimulus category by individual L2/3 neurons of wS1, whereas passive exposure to a repeated tactile stimulation does not. Our results extend growing evidence supporting that perceptual learning is accompanied by enhanced representation of task-relevant features in primary sensory cortex (Corbo et al., 2022; Kato et al., 2015; Poort et al., 2015; Schoups et al., 2001; Schumacher et al., 2022; Xin et al., 2019).

**Figure 3.**
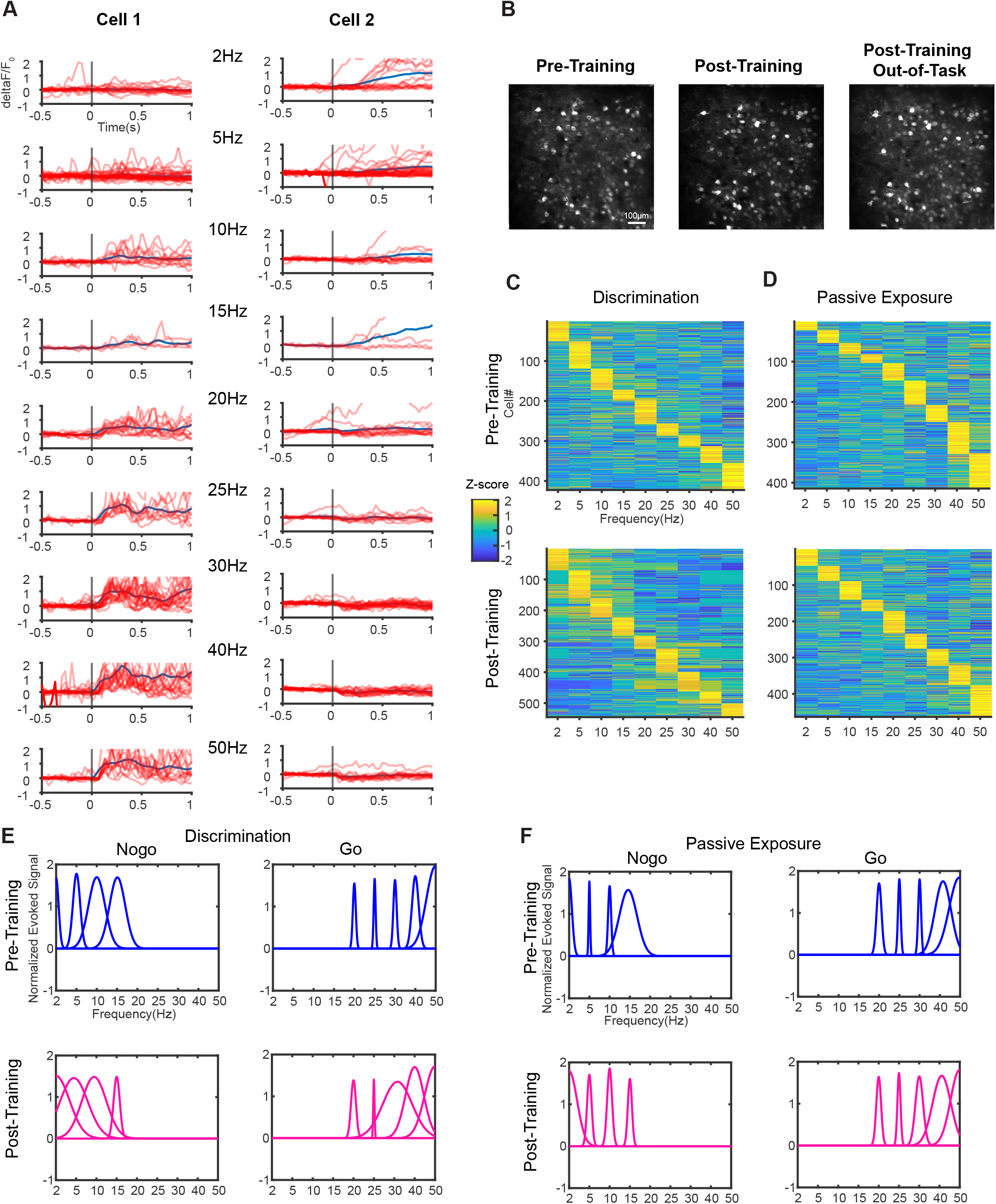
Training in tactile discrimination alters neuronal tuning in L2/3 of wS1. (A) Representative examples of GCaMP7f-expressing cells in L2/3 of wS1 that respond to either Go cues (Cell 1) or NoGo cues (Cell 2). The two cells were present in the same field of view, and their activities recorded in the same session. Red and blue traces indicate raw *ΔF/F_o_* over time from individual trials and their average, respectively. (B) Example field of view showing GCaMP7f-expressing neurons, generated by calculating maximum fluorescence across z-axis of image stacks from 10 randomly selected trials of a pretraining session, post-training session, and post-training out-of-task session. (C) Frequency tuning of individual neurons, before (top) and after (bottom) training in the discrimination task. Color bar indicates z-score of mean evoked *ΔF/F_0_* response, calculated using frames within the first 700 ms after stimulus onset. Neurons in the heatmaps are sorted by ‘preferred frequency’ - i.e., the stimulus frequency associated with peak evoked *ΔF/F_0_* - in the ascending order. Heatmaps were calculated using baseline testing sessions and sessions after the animals became an expert at the frequency discrimination. (D) Same as *C* but for the control group (Passive exposure) that underwent the same procedure as trained mice except that licking did not result in reward or punishment in this group. (E) Frequency tuning of L2/3 population estimated using Gaussian function fitted to the average tuning curve derived from neurons with the same preferred frequency. Mean stimulus-evoked signals were z-scored across all frequencies to generate single neuron tuning curves. (F) Same as *E* but for the passive exposure control.

**Figure 4.**
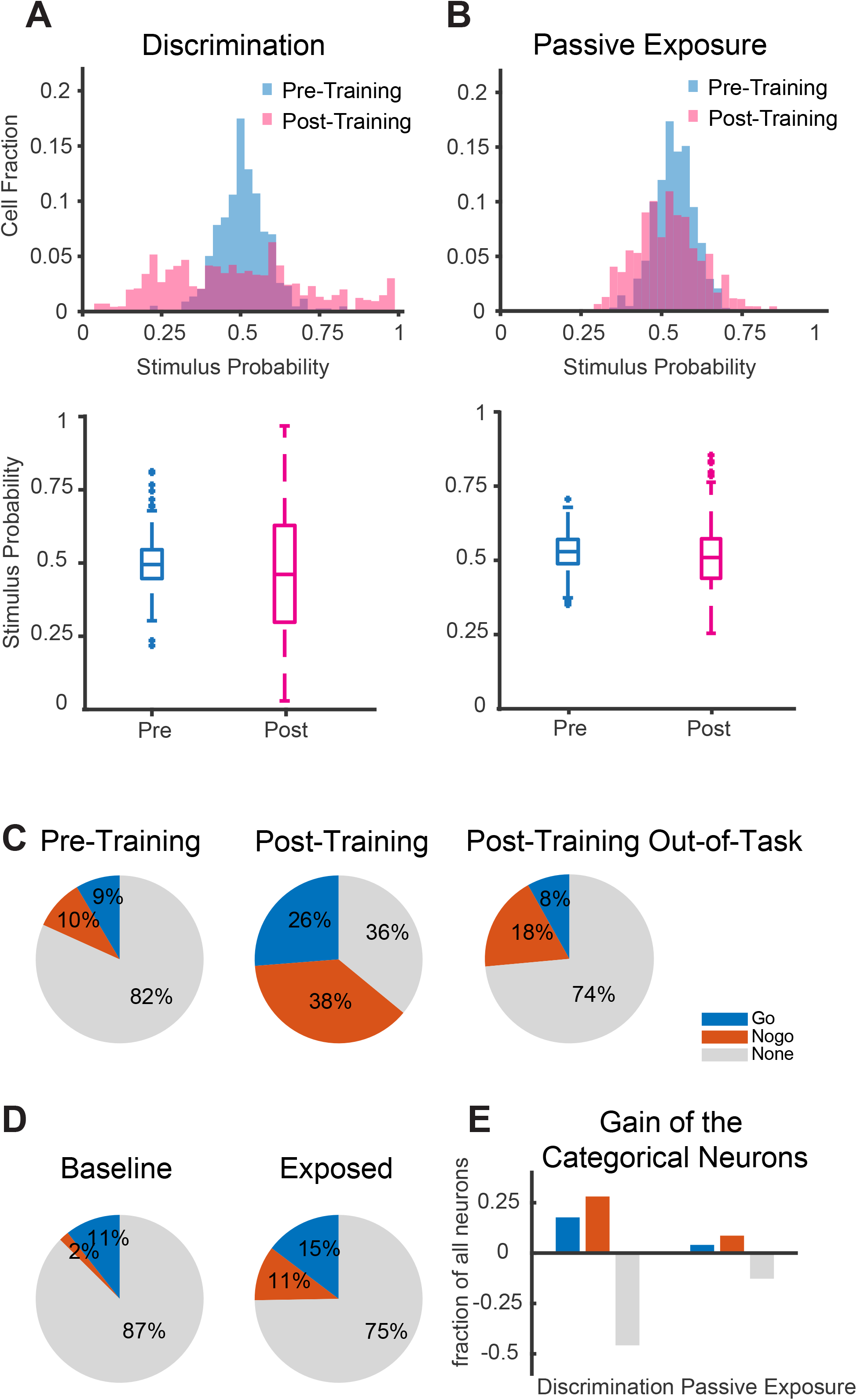
Training in tactile discrimination increases category selectivity of L2/3 neurons in wS1. (A) (top) Histograms showing stimulus probability, a ROC-based metric that quantifies trial-bytrial co-fluctuation of neuronal response (evoked *ΔF/F_0_*) and stimulus category (Go or NoGo). (bottom) Box and whisker plots showing median and interquartile range of stimulus probability values from pre- (blue) and post-training (magenta) sessions. Pre-Training: Median SP = 0.495, 25 percentile SP = 0.447, 75 percentile SP = 0.545, Interquartile Range (IQR) = 0.098. Post-Training: Median SP = 0.459, 25 percentile SP = 0.296, 75 percentile SP = 0.627, IQR = 0.331. (B) Same as *A* but for the passive exposure control. Pre-Exposure: Median SP = 0.528, 25 percentile SP = 0.488, 75 percentile SP = 0.569, IQR = 0.081. Post-Exposure: Median SP = 0.508, 25 percentile SP = 0.439, 75 percentile SP = 0.572, IQR = 0.133. (C) (Left) Pie chart of proportions of ‘Go’ category neurons (blue), ‘NoGo’ category neurons (orange), and non-categorical neurons (grey) in pre-training sessions (n = 6 mice). Cells with 95% CI of stimulus probability (SP) > 0.5 are defined as ‘Go’ category neurons, < 0.5 as ‘NoGo’ category neurons. If 95% CI of SP includes 0.5, they were defined as non-categorical. (Middle) same as left but for post-training sessions (n = 6 mice). (Right) same as left but for post-training out-of-task sessions (n = 4 mice). (D) (Left) same as *C* left but for the passive exposure group during baseline sessions (n = 4 mice). (Right) same as *C* left but for post-passive exposure sessions (n = 4 mice). (E) Change in fractions of ‘Go’, ‘NoGo’, and non-categorical neurons through discrimination training or passive exposure to repeated stimulation.

### Enhanced category selectivity in wS1 neurons is context-dependent

An unanswered question is whether this enhancement of category encoding in wS1 neurons is only evident during discrimination task performance, or persists outside the behavioral paradigm. If the enhanced category selectivity is primarily driven by local synaptic plasticity, it should extend to stimuli presented outside of the task performance. On the other hand, context-dependent enhancement of category selectivity would point to top-down modulation playing a role in the expression of learning-induced changes. To address these issues, we examined the presence of training-induced category selectivity in trained mice behaviorally disengaged from the task. We characterized frequency tuning of L2/3 neurons in a subset of well-trained mice that were satiated for water (4 mice) following the task performance on the same day. We found that the distribution of the peak of activity became comparable to that observed in pre-training sessions (**Figure 5A**). Learning-induced broadening of neuronal tuning curves was not evident at any stimulus frequency (**Figure 5B**). The distribution of stimulus probability was also comparable to that found in naive animals (**Figures 5C and 5D**). Therefore, the learning-induced enhancement in encoding of stimulus category is expressed specifically during task performance.

**Figure 5.**
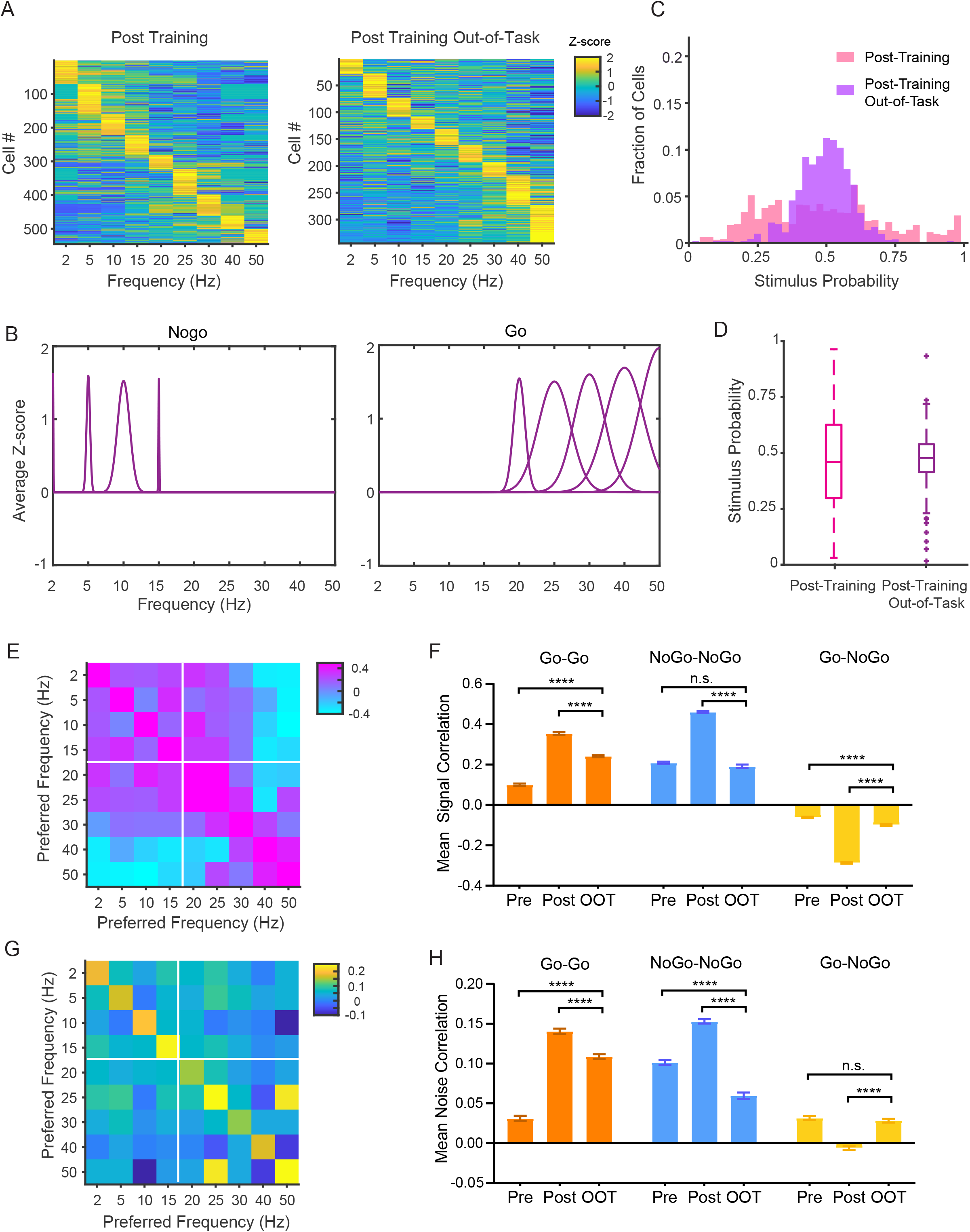
Enhanced category representations are context-dependent. (A) Frequency tuning of individual wS1 L2/3 neurons from post-learning sessions during task performance versus out-of-task. (B) Frequency tuning of L2/3 population. Mean stimulus-evoked signals were z-scored across all frequencies to generate a tuning curve for each neuron. Population tuning curves were then estimated using Gaussian function fitted to the average tuning curve calculated from neurons that share the same preferred frequency. (C) Histogram showing distribution of stimulus probability of individual neurons from post-training in-task (data from **Figure 5F**) and out-of-task conditions. (D) Box and whisker plot showing stimulus probability of individual neurons from post training in-task (data from **Figure 5F**) and out-of-task conditions. Out-of-task: median SP = 0.481, 25 percentile SP = 0.419, 75 percentile SP = 0.543, IQR = 0.124. (E) Color in each bin indicates mean pairwise signal correlation among neurons whose preferred frequency is indicated by x and y coordinate of that bin. Data from post-training out-of-task session in expert animals. (F) Mean signal correlations in expert animals during out-of-task (OOT) sessions. OOT: Go-Go (orange), 0.242 ± 0.006; NoGo-NoGo (blue), 0.192 ± 0.009; Go-NoGo (yellow), −0.099 ± 0.005. Pre-training vs. OOT: t = 16.54, p < 0.0001, df = 11514; post-training vs. OOT: t = 12.67, p < 0.0001, df = 12545; pre-training vs. OOT: t = 1.686, p = 0.0918, df = 6280; post-training vs. OOT: t = 25.82, p < 0.0001, df = 9347; pre-training vs. OOT: t = 5.778, p < 0.0001, df = 15993; post-training vs. OOT: t = 26.78, p < 0.0001, df = 20127. Unpaired t-tests. (G) Same as panel *F* but for noise correlation. (H) Same as *F* but for noise correlation. OOT: Go-Go (orange), 0.109 ± 0.003; NoGo-NoGo (blue), 0.060 ± 0.004; Go-NoGo (yellow), 0.028 ± 0.002. Pre-training vs. OOT: t = 17.35, p < 0.0001, df = 11514; post-training vs. OOT: t = 7.194, p < 0.0001, df = 12545; pre-training vs. OOT: t = 7.722, p < 0.0001, df = 6280; post-training vs. OOT: t = 16.76, p < 0.0001, df = 9347; pre-training vs. OOT: t = 0.9408, p = 0.3468, df = 15993; post-training vs. OOT: t = 9.880, p < 0.0001, df = 20127. Unpaired t-tests.

### Learning increases correlations among neurons based on their stimulus category, in a context-dependent manner

Analyses described thus far focused on the frequency tuning of individual wS1 neurons. Prior studies have demonstrated that perceptual learning can primarily alter noise correlations, relationships of signal and noise correlations, and/or population coding even when there is little effect on neuronal tuning (Gu et al., 2011; Jeanne et al., 2013; Ni et al., 2018). We next tested whether increases in category selectivity of individual neurons are accompanied by network-level reorganization by focusing on neuronal correlations in wS1. Pairwise signal and noise correlations respectively quantify tuning curve similarity and trial-to-trial co-variability of responses (Cohen and Kohn, 2011). Noise correlations are an estimate of synaptic connectivity and/or shared inputs (Ko et al., 2011). By recording large cell populations over a broad range of stimulus frequencies, we were able to estimate ‘correlation matrices’ -– i.e., how signal and noise correlations vary as a function of combinations of frequency preferences in cell pairs. We observed learning-induced enhancement of signal correlations (or frequency tuning similarity) among cell pairs that shared a preferred category. Learning increased signal correlations among pairs that shared a preference toward Go cues (high frequency, **Figures 6A and 6B**) (pre: 0.100 ± 0.006, post: 0.354 ± 0.007), possibly by selectively broadening their tuning toward multiple frequencies associated with reward (**Figure 5D**). Likewise, signal correlations among cell pairs that prefer NoGo cues (low frequency) significantly increased through learning (pre: 0.209 ± 0.006, post: 0.461 ± 0.005). Signal correlations across two pools of neurons that prefer opposite categories (Go-NoGo) became strongly negative (pre: −0.062 ± 0.004, post: −0.287 ± 0.004), suggesting they became more dissimilar (**Figures 6A and 6B**). Although passive exposure resulted in statistically significant changes in signal correlations, the magnitude of change was much smaller compared with what occurred during discrimination task learning. Moreover, Go-Go pairs showed a decrease, rather than an increase, of signal correlations under passive exposure (**Figure 6B**). Therefore, learning enhances tuning similarity in L2/3 neurons in a category-aligned manner.

**Figure 6.**
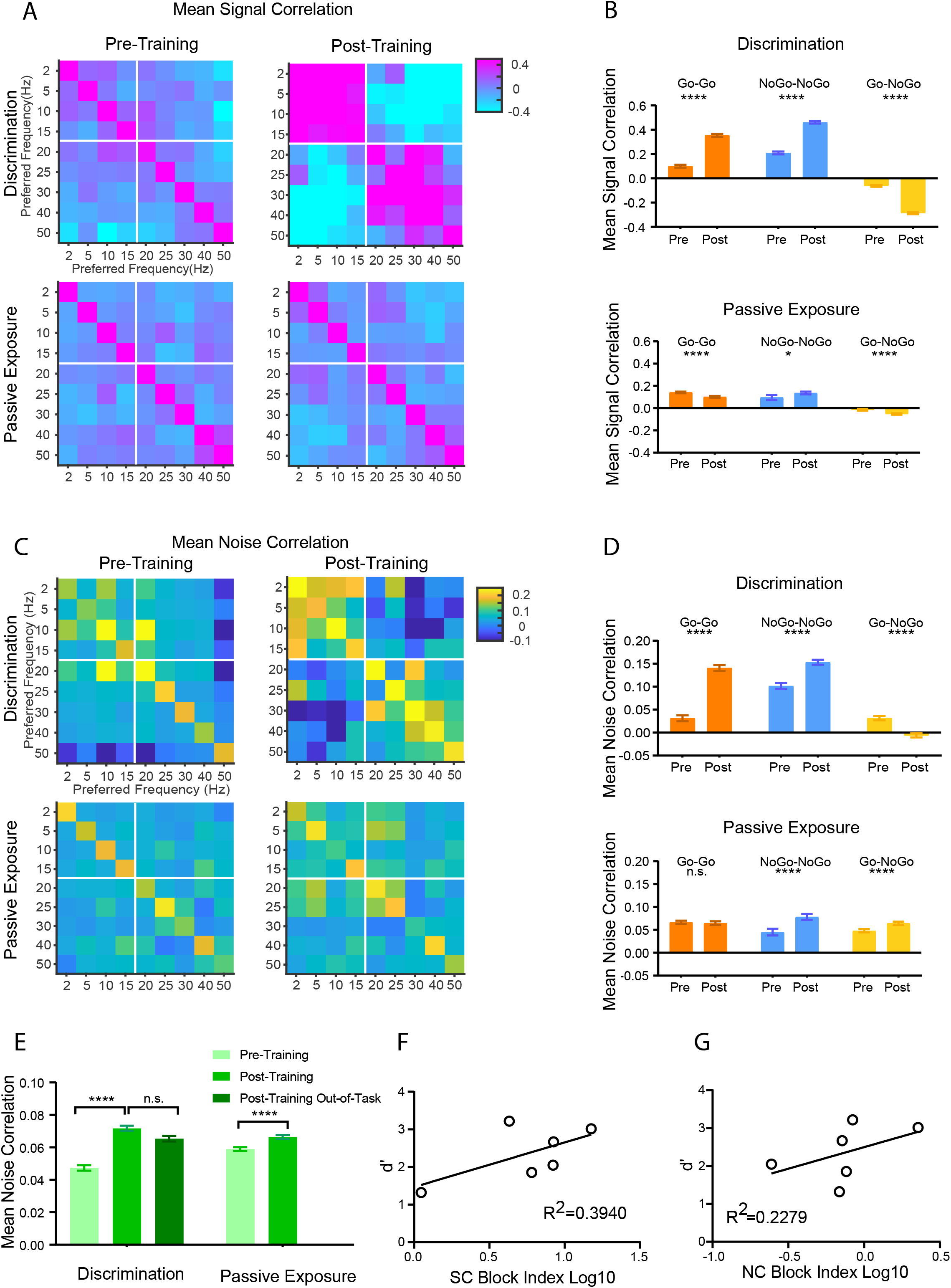
Neurons with shared category preference show an increase in correlations. (A) Color in each bin indicates mean pairwise signal correlation among neurons whose preferred frequency is indicated by *x* and *y* coordinate of that bin. *Left* and *right* panels represent data before and after training. White lines indicate the boundary between Go and NoGo cues. (B) (top) Mean signal correlations among pairs that share the same category preference (‘Go- Go’ or ‘NoGo-NoGo’ pairs) were prominently elevated in expert animals (Post) compared with novice (Pre). Go-Go (orange), pre: 0.100 ± 0.006, post: 0.354 ± 0.007, t=27.75, df = 11483, p < 0.0001; NoGo-NoGo (blue), pre: 0.209 ± 0.006, post: 0.461 ± 0.005, t=31.82, df = 11459, p < 0.0001. Mean signal correlations across neurons that prefer different categories (‘Go-NoGo’ pairs) showed significant decreases after learning. Go-NoGo (yellow), pre: −0.062 ± 0.004, post: −0.287 ± 0.004, t= 35.94, df = 22090, p < 0.0001 (unpaired *t*-test). (bottom) Same as *top* panels but for the control group that received the repeated passive whisker deflection (passive exposure). Mean signal correlations either decreased or slightly increased among pairs that share the same category preference (‘Go-Go’ or ‘NoGo-NoGo’ pairs) after the passive exposure (post) compared with the baseline (pre). Go-Go (orange), pre: 0.142 ± 0.004, post: 0.102 ± 0.004, t=7.445, df = 24772, p < 0.0001; NoGo-NoGo (blue), pre: 0.097 ± 0.011, post: 0.136 ± 0.006, t = 3.091, df = 5329, p = 0.026. Mean signal correlations across neurons that prefer different categories (Go-NoGo) showed modest but significant decreases. Go-NoGo (yellow), pre: −0.014 ± 0.004, post: −0.053 ± 0.003, t = 7.391, df = 21743, p < 0.0001. Unpaired *t*-test for all comparisons. (C) Color in each bin indicates mean pairwise noise correlation among neurons whose preferred frequency is indicated by *x* and *y* coordinate of that bin. *Left* and *right* panels represent data before and after training. White lines indicate the boundary between Go and NoGo cues. (D) (top) Mean noise correlations among pairs that share the same category preference (‘Go- Go’ or ‘NoGo-NoGo’ pairs) were higher in expert animals (post) compared with novice (pre). Go-Go (orange), pre: 0.031 ± 0.003, post: 0.141 ± 0.003, t=23.45, df = 11483, p < 0.0001; NoGo-NoGo (blue), pre: 0.101 ± 0.003, post: 0.153 ± 0.003, t=11.85, df = 11459, p < 0.0001; On the other hand, mean noise correlations decreased across neurons that prefer different categories (Go-NoGo) in expert animals. Go-NoGo (yellow), pre: 0.032 ± 0.002, post: −0.006 ± 0.002, t= 11.19, df = 22090, p < 0.0001. Unpaired *t*-test for all comparisons. (bottom) Same as *top* panels but for the control group that received the repeated passive whisker deflection (passive exposure). Mean noise correlations did not change among pairs that prefer ‘Go’ category but increased among pairs that prefer ‘NoGo’ category, after passive exposure. Go-Go (orange), pre: 0.067 ± 0.002, post: 0.065 ± 0.002, t = 0.805, df = 24772, p = 0.421; NoGo-NoGo (blue), pre: 0.045 ± 0.004, post: 0.078 ± 0.003, t = 5.345, df = 5329, p < 0.0001. Mean noise correlations increased across neurons that prefer different categories after passive exposure. Go-NoGo (yellow), pre: 0.048 ± 0.002, post: 0.064 ± 0.002. t = 5.924, df = 21743, p < 0.0001. Unpaired *t*-test for all comparisons. (E) Mean noise correlations among all neurons. Discrimination: Pre: 0.047 ± 0.002, n = 18403 pairs, Post: 0.072 ± 0.002, n = 26635 pairs, t = 10.20, p < 0.000001, df = 45036; Post Out-of-Task: 0.065 ± 0.002, n = 15390 pairs, (Post vs. Post Out-of-Task) t = 2.545, p =0.010944, df = 42023, (Pre vs. Post Out-of-Task) t = 7.331, p < 0.000001, df = 33791. Passive Exposure: Pre: 0.059 ± 0.001, n = 22380 pairs; Post: 0.066 ± 0.001, n = 29470 pairs, t = 4.194, p = 0.00027, df = 51848. Unpaired t-tests. (F) Performance of individual mice (d’) plotted against signal correlation block index (see Methods). (R^2^ = 0.394; p = 0.182) (G) Same as *F* but showing d’ against noise correlation block index (see Methods). (R^2^ = 0.228; p = 0.338)

Comparison of noise correlation matrices before and after learning revealed a similar trend as signal correlation. We found that learning increased noise correlations for pairs of neurons with more similar category preferences (Go-Go, pre: 0.031 ± 0.003, post: 0.141 ± 0.003; NoGo- NoGo, pre: 0.101 ± 0.003, post: 0.153 ± 0.003) (**Figures 6C and 6D**). On the other hand, cell pairs with opposite preference showed a significant decrease in noise correlations (Go-NoGo, pre: 0.032 ± 0.002, post: −0.006 ± 0.002) (**Figures 6C and 6D**), consistent with prior observations in primate V1 (Bondy et al., 2018). Passive exposure did not alter noise correlation among Go-Go pairs and slightly increased noise correlation among Go-NoGo pairs with opposite category preference as opposed to the observed decrease in the active learning group.

Do the learning-induced changes in neuronal correlations persist outside the context of task performance? To address this question, we calculated signal and noise correlation matrices using trials during the out-of-task condition in which animals were satiated, thus not actively performing the task (**Figures 5E-H)**. Under this condition, ‘within-pool’ signal and noise correlations (Go-Go and NoGo-NoGo) significantly decreased in magnitude compared with levels during active task performance (**Figures 5F and 5H**). ‘Across-pool’ signal and noise correlations (Go-NoGo) also changed during the out-of-task condition to become similar to pretraining levels. Note that the mean noise correlation, which was calculated across all neurons regardless of category preference, slightly increased through active training or passive exposure (**Figure 6E**) and remained elevated during the out-of-task condition. Therefore, the mean noise correlation alone did not differentiate between active learning versus passive exposure or between task engagement and out-of-task contexts.

Next, we asked whether and how changes in pairwise correlations are related to an animal’s behavioral performance. To quantify relative strengths of ‘within-pool’ versus ‘across-pool’ correlations in individual animals, we devised a metric called ‘correlation block index’ (see Methods). Both signal and noise correlation block indices tended to be greater in animals with better behavioral performance (signal correlation, R^2^ = 0.394; noise correlation, R^2^ = 0.228), although this observation did not reach statistical significance (**Figures 6F and 6G**). Taken together, we conclude that learning increases ‘within-pool’ correlations while decreasing ‘across-pool’ correlations. Critically, this learning-induced re-organization of neuronal correlations is behavioral context-dependent.

### Learning enhances coupling between signal and noise correlations in wS1 neurons

Prior studies suggest that perceptual learning changes the relationship of signal and noise correlations in a direction that reduces noise correlation among highly signal-correlated pairs (Gu et al., 2011; Jeanne et al., 2013). We asked if the observed learning-induced changes in correlation matrices (**Figure 6**) reflect an altered relationship at the level of individual pairs of neurons. We divided neuronal pairs into subgroups of Go-Go, NoGo-NoGo and Go-NoGo, depending on the preferred category of individual neurons. Consistent with previous studies (Bondy et al., 2018; Cohen and Kohn, 2011; Kwon et al., 2018), there was a significant positive relationship between signal and noise correlations even before training (**Figure 7A**). Note that signal and noise correlations are not mathematically related, because noise correlation quantifies trial-by-trial co-fluctuation around the mean evoked response for each stimulus frequency. Noise correlations grew in magnitude through learning among strongly signal-correlated pairs in the ‘within-pool’ subgroups (Go-Go and NoGo-NoGo) (**Figures 7A and 7B**). On the other hand, strong negative noise correlations emerged among negative signal-correlated pairs in the ‘across-pool’ subgroup (Go-NoGo) (**Figure 7C**). The enhanced correlation between neurons of shared category preference is consistent with formation of ‘like-to-like’ local subnetworks among L2/3 neurons, as shown by prior theoretical (Ocker and Doiron, 2019) and experimental works (Khan et al., 2018; Ko et al., 2011; Najafi et al., 2020). Together, these results demonstrate that tactile discrimination learning drives the emergence of L2/3 neuronal subnetworks that encode distinct stimulus categories.

**Figure 7.**
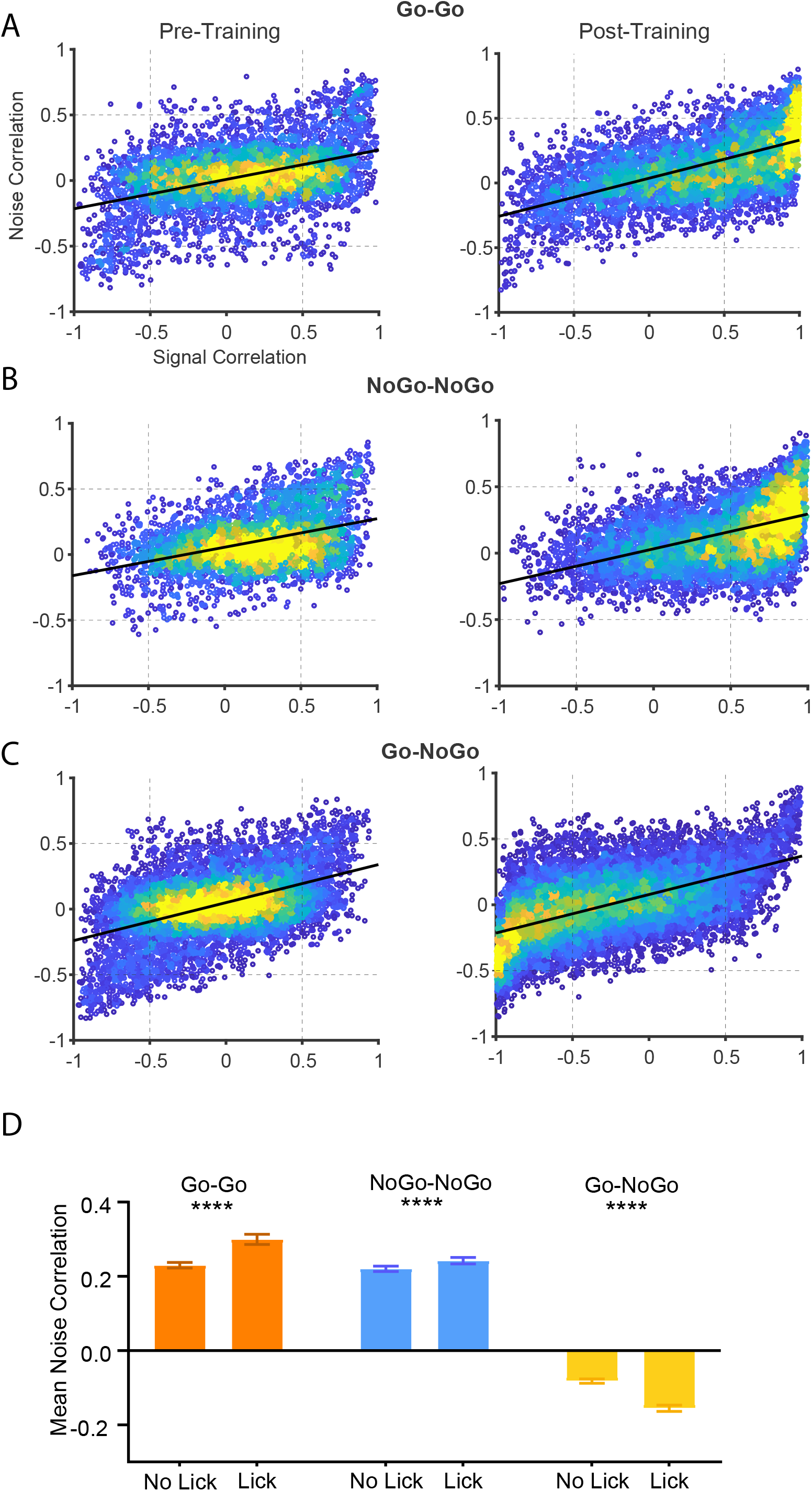
Noise correlations among signal-correlated neurons increase through learning. (A) Noise correlation is plotted against signal correlation for neuron pairs that preferentially respond to a Go stimulus frequency. Plots for pre-training (Left, correlation coefficient = 0.421) and post-training (Right, correlation coefficient = 0.594) sessions are shown for comparison. (B) Same as *A* but for pairs that prefer a Nogo stimulus frequency (Pre: correlation coefficient = 0.393; Post: correlation coefficient = 0.475). (C) Same as *A* but between neurons that prefer different stimulus categories (Pre: correlation coefficient = 0.480; Post: correlation coefficient = 0.576). (D) Mean noise correlation on lick versus no lick trials. ****, p < 0.0001 for all comparisons (paired t test). Go-Go (orange), No-Lick: 0.230 ± 0.004, Lick: 0.300 ± 0.007, t=-11.060, df = 2455, p < 0.0001; NoGo-NoGo (blue), No-Lick: 0.221 ± 0.004, Lick: 0.242 ± 0.004, t=-5.7031, df = 4293, p < 0.0001; Go-NoGo (yellow), No-Lick: −0.082 ± 0.003, Lick: −0.156 ± 0.004, t= 19.181, df = 5630, p < 0.0001 (paired *t*-test).

Does the learning-induced reorganization of noise correlations have a functional significance? To address this question, we sub-selected trials based on the animal’s behavioral responses and asked if mean noise correlation varied on trials with different responses. Category-specific neurons were defined based on stimulus probability (SP) values (**Figure 4A**); a neuron whose mean SP ± 95% CI > 0.5 or < 0.5 was respectively defined as a Go or NoGo category-specific neuron (Kwon et al., 2016). Within-pool noise correlations (Go-Go and NoGo-NoGo) increased in magnitude, whereas across-pool noise correlations (Go-NoGo) became more negative on trials in which the animal generated the learned behavioral response (Go-Go: no-lick, 0.230 ± 0.004; lick, 0.300 ± 0.007; NoGo-NoGo: no-lick, 0.221 ± 0.004; lick, 0.242 ± 0.004; Go-NoGo: no-lick, −0.082 ± 0.003; lick, −0.156 ± 0.004 (**Figure 7D**). Based on these results, we conclude that learning-induced L2/3 subnetworks of correlated neurons facilitate the conversion of sensory input into learned behavioral responses.

### Noise correlation impacts representation of stimulus category by wS1 population activity

Prior theoretical studies have established that neuronal correlations affect the information content in a population of neurons (Abbott and Dayan, 1999; Panzeri et al., 2022). To understand how learning-induced changes in neuronal correlations impact the population code, we constructed a support vector machine (SVM)-based classifier to decode stimulus categories from L2/3 population activity (**Figure 8A**). The decoding accuracy was used as a proxy for ‘information’ represented by the neuronal population. There was a modest increase in decoding accuracy after passive exposure from 0.605 ± 0.021 to 0.695 ± 0.053, but it was not statistically significant (p = 0.150; paired *t*-test). On the other hand, learning significantly improved decoding accuracy from 0.652 ± 0.030 to 0.798 ± 0.035 (p = 0.002; paired *t*-test), which closely matched the animal’s classification accuracy (mean = 0.836; n = 6) (**Figure 8B**). To test the impact of noise correlation on population decoding, we selectively removed the noise correlation by shuffling the trial labels separately for each neuron (**Figure 8C**). Shuffling was nested within each frequency, so the frequency tuning of individual neurons was not altered by this procedure (data not shown). We repeated shuffling for 100 iterations and calculated the decoding accuracy after each iteration. We then asked if the decoding accuracy calculated using pre-shuffled data fell below or went above the mid-95% interval of the 100 post-shuffle accuracy values. In 5 of 6 expert mice trained to perform the discrimination task, shuffling out noise correlation significantly improved decoding accuracy (Original: 0.820 ± 0.038; Shuffled: 0.884 ± 0.045) (**Figure 8E; Table 1**). On the other hand, shuffling out the noise correlation did not change or slightly decreased the stimulus decoding accuracy in other task conditions including pre-training and passive exposure (**Figure 8E**). These results show that the changes in noise correlation associated with learning have a detrimental effect on the encoding of the task-related stimulus category in wS1 population activity, consistent with prior observations (Ni et al., 2018; Valente et al., 2021). Next, we assessed how learning-induced changes in noise correlation impact the decoding of an animal’s behavioral choice (to lick or not). To our surprise, shuffling out the noise correlation had mixed effects with no overall change in choice decoding accuracy (Original: 0.827 ± 0.035; Shuffled: 0.827 ± 0.038) (**Figure 8F; Table 2**). Therefore, learning-induced changes in noise correlation do not impact decoding of an animal’s choice from population activity, although they reduce the amount of stimulus information in wS1 L2/3. Together with the results showing that noise correlations are elevated on lick trials (**Figure 7D**), our results suggest that correlations facilitate the transformation of sensory input into learned behavioral responses despite their detrimental effects on sensory encoding.

**Figure 8.**
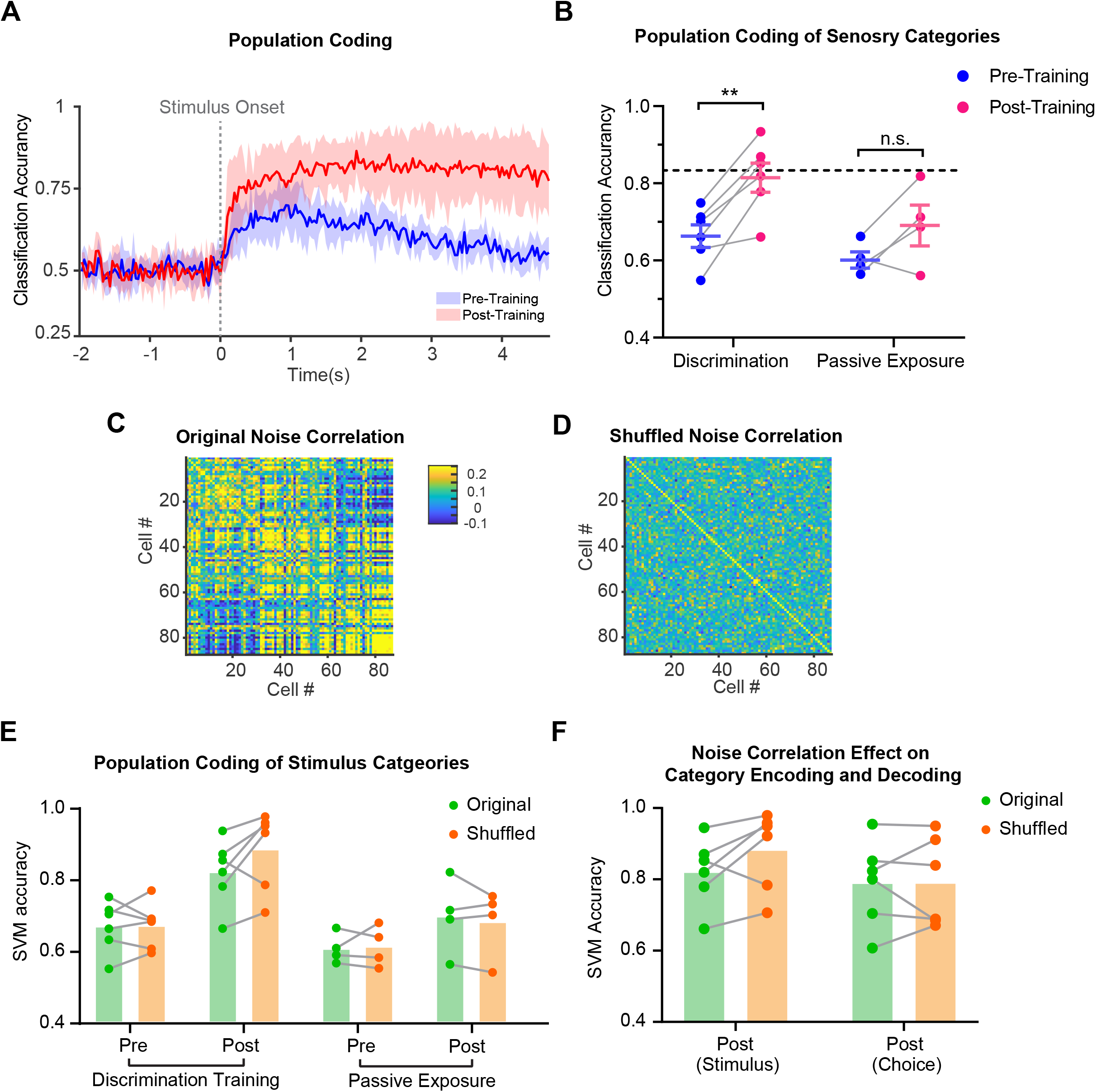
Impact of pairwise correlation on population decoding of stimulus and choice. (A) Performance of a SVM classifier (mean ± SEM across mice) in decoding stimulus category from population activity at each time point of image acquisition. (B) Classification performance reached higher levels in both discrimination learning and passive exposure groups, although the increase was not statistically significant in the passive exposure group. Each dot indicates an animal. Discrimination: pre, 0.667 ± 0.029; post, 0.820 ± 0.021; t = 5.308, df = 5, p = 0.003. Passive exposure: pre, 0.605 ± 0.021; post, 0.695 ± 0.053; t = 1.923, df = 3, p = 0.150. Dashed line indicates expert animal’s average performance (0.836, n = 6). Two tailed paired *t-*tests for all comparisons. (C) Procedure for removing pairwise noise correlation without affecting frequency tuning curves. (top) Example heatmaps showing frequency tuning of individual neurons before and after shuffling trial labels. (bottom) Pairwise noise correlation matrix before and after shuffling trial labels. (D) Performance of a SVM classifier in decoding stimulus category from population activity before and after removing pairwise noise correlation (see Methods). Each dot represents an average Decoding accuracy calculated from 100 iterations normalized to the original Decoding accuracy for each animal. 5 of 6 late-lickers trained to perform the discrimination task showed statistically significant improvement in Decoding accuracy when noise correlation was removed. On average, decoding accuracy improved from 0.820 ± 0.038 to 0.884 ± 0.045 in post-training group. In all other conditions, the effects were mixed. (E) Comparing impact of removing pairwise noise correlation on decoding stimulus category versus animal’s choice from population activity acquired in expert animals (Original: 0.827 ± 0.035; Shuffled: 0.827 ± 0.038; n = 6 mice).

**Table 1.** Accuracy of decoding stimulus category using wS1 population activity before and after discrimination training in the presence or absence of noise correlations. Each row indicates individual mouse. Decoding accuracy for shuffled data was averaged across 100 iterations. The mean accuracy and 95 % confidence interval are shown.

| Decoding Accuracy (Stimulus) |  |  |  |  |  |
| --- | --- | --- | --- | --- | --- |
| Pre-Training |  |  | Post-Training |  |  |
| Original | Shuffled | 95 CI of shuffled | Original | Shuffled | 95 CI of shuffled |
| 0.702 | 0.768 | [0.716,0.808] | 0.820 | 0.958 | [0.916,0.988] |
| 0.713 | 0.683 | [0.633,0.735] | 0.870 | 0.949 | [0.911,0.974] |
| 0.630 | 0.604 | [0.551,0.647] | 0.661 | 0.706 | [0.639,0.783] |
| 0.548 | 0.593 | [0.541,0.637] | 0.779 | 0.779 | [0.888,0.963] |
| 0.750 | 0.689 | [0.644,0.7330] | 0.935 | 0.975 | [0.955,0.990] |
| 0.661 | 0.681 | [0.646,0.716] | 0.853 | 0.784 | [0.758,0.811] |

**Table 2.** Accuracy of decoding stimulus category or animal’s choice using wS1 population activity in trained animals. Each row indicates individual mouse. Decoding accuracy for shuffled data was averaged across 100 iterations. The mean accuracy and 95% confidence interval are shown.

| Decoding Accuracy (Post-Training) |  |  |  |  |  |
| --- | --- | --- | --- | --- | --- |
| Stimulus |  |  | Choice |  |  |
| Original | Shuffled | 95 CI of shuffled | Original | Shuffled | 95 CI of shuffled |
| 0.702 | 0.768 | [0.716,0.808] | 0.607 | 0.671 | [0.598,0.740] |
| 0.713 | 0.683 | [0.633,0.735] | 0.852 | 0.839 | [0.800,0.878] |
| 0.630 | 0.604 | [0.551,0.647] | 0.704 | 0.689 | [0.613,0.758] |
| 0.548 | 0.593 | [0.541,0.637] | 0.824 | 0.911 | [0.867,0.949] |
| 0.750 | 0.689 | [0.644,0.7330] | 0.955 | 0.950 | [0.920,0.975] |
| 0.661 | 0.681 | [0.646,0.716] | 0.800 | 0.685 | [0.633,0.751] |

## Discussion

In this study, we demonstrate that training in a wS1-dependent tactile frequency discrimination task results in enhanced representation of task-related (i.e., discriminated) categories in mouse wS1 layer 2/3 neurons. Neurons gained the ability to represent stimulus categories by broadening their frequency tuning curves. Critically, this learning-induced plasticity was not observed in animals passively experiencing repetitive whisker stimulation, or when actively trained animals were disengaged from the task performance. Task learning increased ‘within-pool’ neuronal correlations while decreased ‘across-pool’ correlations such that neurons became increasingly aligned along the direction of stimulus category, indicating increased ‘like-to-like’ interactions. The learning-induced re-organization of noise correlations had detrimental effects on the amount of sensory information estimated using population decoding accuracy, but it did not constrain an animal’s decision during task performance. Furthermore, ‘like-to-like’ interactions were elevated on trials where trained animals exhibited the decision-related behavior. This indicates that ‘like-to-like’ interactions may facilitate propagation of sensory information to downstream areas that execute learned behavioral responses, although they simultaneously limit the amount of sensory information within wS1. Taken together, we showed that representation of task-related categories emerges through changes in selectivity of individual neurons and in the structure of neuronal correlations even at the very first stages of cortical sensory processing. These learning-induced changes likely facilitate propagation of sensory information from primary sensory cortex to downstream areas.

Neurons in the adult mouse primary sensory cortex undergo various forms of plastic changes during learning. Extensive studies in mouse V1 L2/3 have documented an increased selectivity for rewarded stimuli, increased number of responsive neurons, reduced trial-to-trial variability of responses, enhanced encoding of non-sensory task-related variables and various changes in tuning curves (Goltstein et al., 2013; Goltstein et al., 2018; Henschke et al., 2020; Jurjut et al., 2017; Poort et al., 2015). These modifications generally result in enhanced discrimination of target versus foil stimuli and correlate with behavioral performance. Our results extend these findings and demonstrate that wS1 neurons also gain selectivity to task-related categories through learning. A main finding of our study is that cortical plasticity during perceptual learning involves changes in pairwise correlations that become increasingly aligned to task-related categories, indicating formation of task-related neuronal subnetworks in the L2/3 of sensory cortex.

Prior studies suggest that these subnetworks of correlated neurons may play an important role in amplifying relevant stimulus features (Peron et al., 2020), reducing dimensionality of sensory information (Nassar et al., 2021) and enhancing propagation and read-out of sensory information (Valente et al., 2021; Zylberberg et al., 2017). A recent work in mouse posterior parietal cortex demonstrated that correlations can benefit task performance even if they decrease sensory information as they enhance the conversion of sensory information into behavioral choices (e.g. ref 22). A novel finding of our study is that this functional role of noise correlations is sharpened by learning. This reveals a novel neural mechanism of learning that adds significantly to the previous seminal work on noise correlations and learning (Cohen and Maunsell, 2009; Ni et al., 2018).

The causal role of subnetworks of correlated neurons in early sensory cortex remains to be addressed by manipulating subnetworks in sensory cortex and assessing the impact on downstream areas in the same brain. A recent study has found that an ensemble of neurons in L2/3 of wS1 play a causal role in driving an animal’s decision-making during a texture discrimination task (Buetfering et al., 2022). The questions of whether and how L2/3 subnetworks propagate sensory information to downstream areas during decision-making may be tackled using simultaneous optogenetic manipulation and calcium imaging.

Synaptic mechanisms underlying learning-induced category-specific increases in noise correlations remain unclear. A potential source of noise correlations includes local synaptic connectivity (Ko et al., 2014), shared afferent input (Shadlen and Newsome, 1998), feedforward (Kanitscheider et al., 2015) and top-down feedback signals (Bondy et al., 2018). In our study, category-specific increases in signal and noise correlations were only observed during active task performance, which points to top-down feedback as a major source of modulation of correlations. Further studies would be required to dissect contributions of different sources. Neuronal subtypes that participate in the subnetwork of correlated neurons need to be determined. Recent studies suggest that inhibitory interneurons as well as excitatory neurons participate in cortical subnetworks (Agetsuma et al., 2018; Khan et al., 2018; Najafi et al., 2020; Wilmes and Clopath, 2019). Excitatory neurons that project to specific downstream areas may be preferentially recruited to the subnetworks (Chen et al., 2015; Kwon et al., 2016; Yamashita and Petersen, 2016). The precise function of different interneuron and projection neuron subtypes in the formation and maintenance of task-relevant subnetworks needs to be assessed in future studies, using cell type-specific imaging and manipulation methods.

## Acknowledgments

This research is funded by Simons Foundation Autism Research Initiative (SFARI) Grant: Sung Eun Kwon 479737. We thank all current and previous members of the Kwon lab for their kind support and suggestions. We also thank Dr. Sara Aton, Dr. Victoria Booth, and Dr. Nao Uchida for providing feedback on this manuscript.

